# Physiological JNK3 Concentrations Are Higher in Motor-related and Disease-implicated Brain Regions of C57BL6/J Mice

**DOI:** 10.1101/2024.01.17.575386

**Authors:** Victoria Godieva, Ferass Sammoura, Sebastian Verrier Paz, Yoonhee Han, Valentina Di Guida, Michael J. Rishel, Jason R. Richardson, Jeremy W. Chambers

## Abstract

The c-Jun N-terminal kinase 3 (JNK3) is a stress-responsive protein kinase primarily expressed in the central nervous system (CNS). JNK3 exhibits nuanced neurological activities, such as roles in behavior, circadian rhythms, and neurotransmission, but JNK3 is also implicated in cell death and neurodegeneration. Despite the critical role of JNK3 in neurophysiology and pathology, its localization in the brain is not fully understood due to a paucity of tools to distinguish JNK3 from other isoforms. While previous functional and histological studies suggest locales for JNK3 in the CNS, a comprehensive and higher resolution of JNK3 distribution and abundance remained elusive. Here, we sought to define the anatomical and cellular distribution of JNK3 in adult mouse brains. Data reveal the highest levels of JNK3 and pJNK3 were found in the cortex and the hippocampus. JNK3 possessed neuron-type selectivity as JNK3 was present in GABAergic, cholinergic, and dopaminergic neurons, but was not detectable in VGLUT-1-positive glutamatergic neurons and astrocytes *in vivo*. Intriguingly, higher JNK3 signals were found in motor neurons and relevant nuclei in the cortex, basal ganglia, brainstem, and spinal cord. While JNK3 was primarily observed in the cytosol of neurons in the cortex and the hippocampus, JNK3 appeared commonly within the nucleus in the brainstem. These distinctions suggest the potential for significant differences between JNK3 actions in distinct brain regions and cell types. Our results provide a significant improvement over previous reports of JNK3 spatial organization in the adult CNS and support continued investigation of JNK3’s role in neurophysiology and pathophysiology.

## Introduction

The c-Jun N-terminal kinases (JNKs) are vital members of the mitogen-activated kinase (MAPK) family and play crucial roles within the central nervous system (CNS). These include but are not limited to, cell differentiation and proliferation, axonal growth and transport, and neuronal cell death [1–5]. The JNK family comprises three distinct isoforms JNK1, JNK2, and JNK3, which are further distinguished from one another by more than ten unique splice variants [6]. While JNK1 and JNK2 are ubiquitously expressed across all tissue types [5], JNK3 is primarily localized in the brain, with minor traces found in the testis and heart [7–9]. Intriguingly, whole-body gene ablation of *Jnk3* in mice only appears to confer neuroprotection to stress-induced dysfunction and death [10–13]. To gain insights into why JNK3 is readily present in the CNS, but only seemingly responsive to stress, a more descriptive anatomical description of JNK3 distribution within the CNS may be required.

To date, the primary narrative regarding JNK3 activities, as with other JNK isoforms, has focused on interrogating its role in neuron death and in the pathogenesis of neurodegenerative diseases [1–3, 14, 15]. Indeed, early studies into JNK3 selective events in the CNS, were based on the generation of *Jnk3^-/-^*mice that had no distinguishable phenotype from wild-type mice except when challenged by kainic acid [10]. The kainic acid-exposed *Jnk3^-/-^* mice exhibited reduced hippocampal neuron loss than their wild-type counterparts [10], thus implicating JNK3 in neuronal apoptosis. The association of JNK3 and neuronal apoptosis was strengthened through a series of studies on cell lines and cultured neurons identifying JNK3 activation as an early cell death response in neurotrophic withdrawal [9, 16], arsenite exposure [17, 18], and other neurologically relevant stressors [19–22]. Thus, it was not surprising when elevated phospho-JNK levels were found in the post-mortem brains of patients diagnosed with neurodegenerative diseases [23–25].

Recent preclinical studies in Alzheimer’s disease (AD) and Parkinson’s disease (PD) models illustrate that induction of JNK3-related activities may lead to neuronal cell death [11, 23, 24, 26–32]. In AD, JNK3 has been shown to phosphorylate the microtubule-associated protein, tau, which is hyperphosphorylated in AD leading to the formation of neurofibrillary tangles [33–35]. Additionally, isoform-selective inhibition of JNK3 in the 3xTg-AD mouse model resulted in reduced tau phosphorylation and improved cognitive function [36]. In various models of neurodegenerative diseases, JNK3-selective inhibitors protect neurons against neurotoxin insults and genetic induction of symptoms [32, 37–39]. Consequently, sustained JNK3 activity may lead to neuron loss (reviewed in [26] and [5]).

The activities of JNK3 in neuronal physiology continue to emerge. Specifically, JNK3 may contribute to neuronal morphology, as JNK3 overexpression in nerve growth factor (NGF)-treated pheochromocytoma (PC12) cells leads to enhanced dendrite sprouting [40]. Complementary to dendritic architecture maintenance, JNK3 may play crucial roles in synaptic neurotransmission, as JNK3 is involved in the assembly and disassembly of microtubules by phosphorylating SCG10, a neuronal growth-associated protein [15]; additionally, JNK3 has been found on vesicular structures in axons [41], wherein, it may engage in the phosphoregulation of dynein and kinesin [41–43]. Accordingly, early studies into JNK3 subcellular localization found the protein on vesicles [44] placing it in close proximity to these motor proteins. Potential neurological contributions of JNK3 have also been described in knockout mice. *Jnk3^-/-^* mice were found to have diminished neurogenesis in adult mice evidenced by a decrease in cells positive for neural precursor markers in the dentate gyrus [45]. Additionally, *Jnk3^-/-^* mice exhibited behavioral phenotypes in a battery of tests [46]; specifically, *Jnk3^-/-^*mice had a lower distance moved and decreased center crossing in the elevated plus maze compared to control animals, while in the Morris water maze, *Jnk3^-/-^* mice trended toward longer training times, increased times in the border zone, and increased total times than control animals [46]. Recent studies now suggest a putative role for JNK3 in metabolic regulation and homeostasis broadening the impact of JNK3 in the CNS [47]. Taken together, these results indicate the possibility of nominal JNK3 activities supporting basal neurological functions.

Although these findings underscore the significance of JNK3 within the brain, a comprehensive understanding of JNK3’s precise localization remains elusive. Initial studies employing the p493F12 antibody for immunohistochemical staining in AD human brains along with *in situ* hybridization revealed that JNK3 is prominently expressed in the cortex, hippocampus, subcortical regions, and the brainstem, with a weaker signal detected in the spinal cord. This study further suggested that the localization of JNK3 was primarily neuronal [8], but lacked the context of a healthy brain. Subsequent research using *in situ* hybridization on adult male Sprague-Dawley rats illustrated high JNK3 expression in the cerebral cortex and hippocampus, with a lesser degree in the striatum. In the hippocampus, high JNK3 expression was reported predominantly within neuron-like cell bodies in CA1, CA3, the subiculum, and the dentate gyrus, while moderate staining was observed in the habenula [48]. Most recently, radiolabeled-aminopyrazoles were developed as isoform-selective probes for activated JNK3 (pJNK3), which were tested for selectivity in a Parkinson’s disease mouse model (Mutator/Park2^-/-^ mice) [49]. These probes found more pJNK3 in the hippocampus and the cortex of the C57BL6/J mice with other regions exhibiting relatively low levels. However, when compared to the 9-month-old Mutator/Park2^-/-^ mice, higher pJNK3 concentrations were detected in the ventral midbrain, suggesting that disease progression may correlate with pJNK3 levels in the ventral midbrain. Despite these advances, basal JNK3 localization in a healthy adult mouse brain remains to be extensively explored with validated JNK3-selective reagents with areas like the brainstem and the spinal cord yet to be thoroughly examined.

The primary aim of this study is to identify JNK3 localization within healthy adult male and female mouse brains. Using a knockout-validated, isoform selective antibody, we report the immunohistochemical and immunofluorescent localization of JNK3 in various regions of the brain and cervical spinal cord and compare histological protein levels to pJNK3/JNK3 levels in dissected brain regions. Furthermore, we identify JNK3-containing neuronal cell types, thereby contributing to a more comprehensive understanding of JNK3’s nature in the healthy adult brain.

## Experimental Procedures

### Materials

General laboratory reagents and cell culture reagents were purchased from ThermoFisher Scientific (Waltham, MA). Immunohistochemical reagents were purchased from Vector Labs, Inc. (Newark, CA). Western blotting reagents were purchased from Bio-Rad Laboratories (Hercules, CA) and Li-COR, Inc. (Lincoln, NE).

### Animal care

All animal care and experimental procedures were performed in compliance with the Institutional Animal Care and Use Committee (IACUC) guidelines at Florida International University. The subjects of the study were 3-month-old C57BL6/J mice (n=4/sex) and 12-month-old C57BL6/J mice, both males (n=9) and females (n=10). The mice underwent two types of procedures. For some mice, transcardial perfusion with formalin was performed for tissue fixation. Briefly, mice were deeply anesthetized with isoflurane and then transcardially perfused with phosphate-buffered saline (PBS), followed by 10% formalinin. The brains were carefully dissected out, post-fixed in 10% formalin for one week and stored in 30% sucrose solution containing 0.1% sodium azide until sectioning on microtome Brains were either sectioned sagittaly or coronally on the Thermo Scientific Microm HM 450 sliding microtome (Walldorf, Germany) into 30 um sections and stored in 96 well plates containing anti-freeze solution. For the other mice, brain regions were dissected for fresh freezing. Mice were euthanized following IACUC-approved procedures. Cerebellum, brainstem, hippocampus, midbrain, striatum, and cortex were quickly and carefully dissected. The brain regions were immediately frozen in dry ice or liquid nitrogen. The frozen tissue samples were then stored at -80°C until homogenizing.

### Immunohistochemical (IHC) Staining

Tissues were thoroughly washed six times with phosphate-buffered saline (PBS) (Gibco #21600-010) for 5 minutes. Next, the tissues were treated with a solution containing 2.5% hydrogen peroxide (30%), 75% methanol, and 22.5% deionized water for 20 minutes on an orbital shaker. Afterward, the tissues were blocked using a PBS solution containing 15% bovine serum albumin (BSA) (Sigma-Aldrich # A7030), 4% normal goat serum (Vector #S-1000), and 0.6% Triton X-100 which was allowed to shake for an hour on an orbital shaker. Subsequently, the primary antibody (Table 1) was diluted with PBS and added to the tissue sections for overnight incubation at 4°C on orbital shaker. Following incubation, tissue sections were washed five times with PBS for 5 minutes. The goat anti-rabbit (Vector Laboratories #BA-1000), or goat anti-mouse (Vector Laboratories #BA-9200) secondary antibodies, conjugated to horseradish peroxidase (HRP), was then added and incubated for 1 hour at room temperature on an orbital shaker. After incubation, tissue sections were washed another five times with PBS for 5 minutes. The sections were then incubated with an avidin-biotin complex (ABC) solution (Vector Laboratories #PK-4000) which was prepared as suggested by the manufacturer for 1 hour at room temperature on an orbital shaker. Sections were washed five times with PBS for 5 minutes. Visualization of the target protein was achieved by adding 3,3’-diaminobenzidine (DAB) solution (Vector Laboratories #SK-4100), and the reaction was stopped by placing the sections in PBS. Afterward, the tissue sections were mounted on glass slides and air-dried. The sections were then rehydrated through a series of ethanol washes (30%,50%,75%,95%, and 100%) and cleared using CitriSolv (Decon Laboratories # 1601). Finally, the tissue sections were coverslipped using DPX mounting medium (Electron Microscopy Sciences #13512), and the slides were allowed to dry completely before microscopic examination on BZ-X All-in-One Fluorescence microscope (Keyence, Itasca, IL).

**Table 1.**
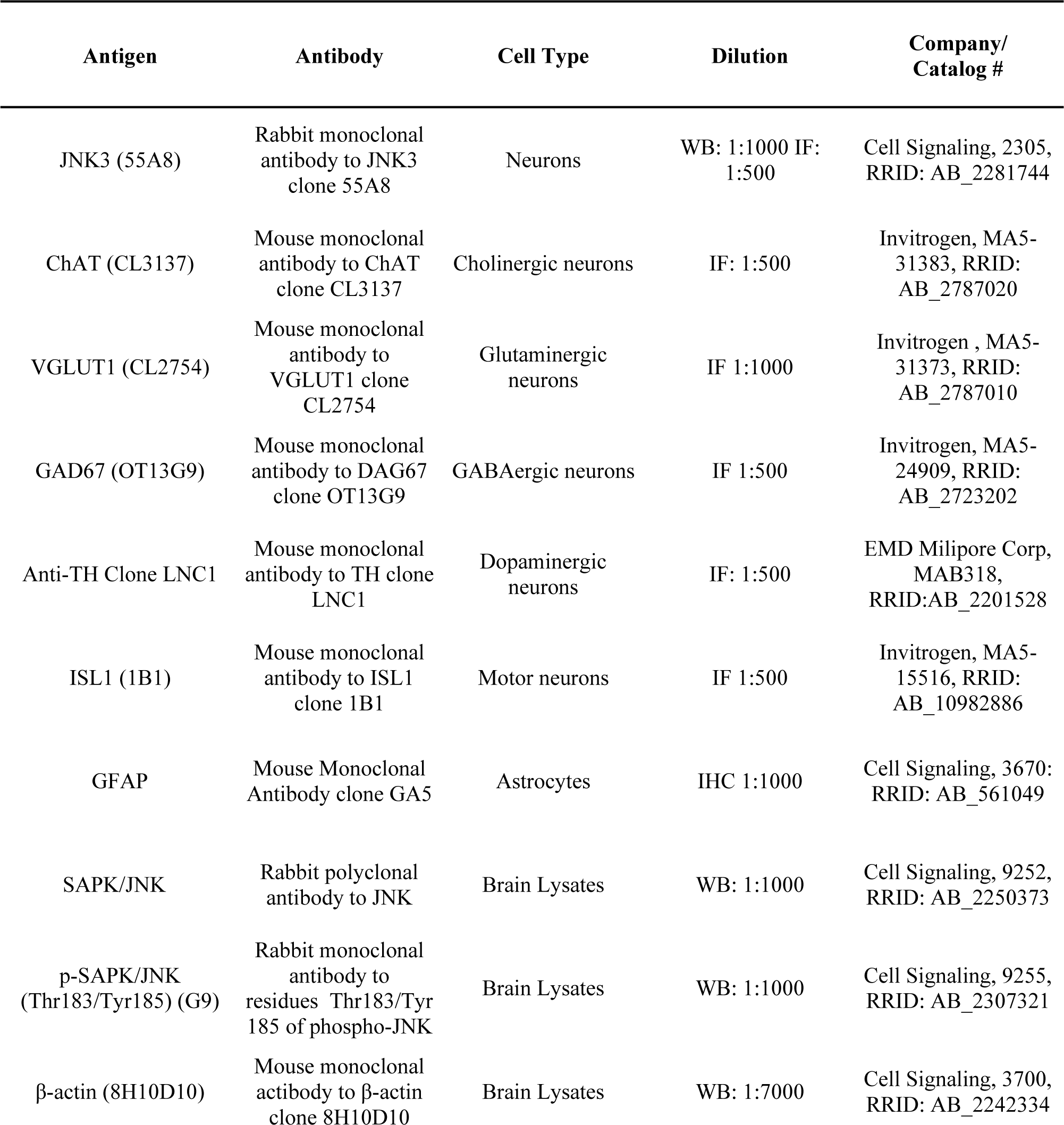
Summary of Antibodies.

### Immunofluorescent (IF) Staining

Tissues were washed overnight with PBS at 4°C shaking on the orbital shaker. Next day, tissues were then permeabilized with PBS solution containing 0.5% Triton X-100 for 20 minutes on an orbital shaker. Following permeabilization, tissues were washed three times with PBS for 10 minutes each. Then tissues were blocked using a solution containing PBS with 10% bovine serum albumin (BSA) and 0.1% Triton X-100 for 1 hour at room temperature shaking on an orbital shaker. Primary antibodies (Table 1), diluted in PBS with 1% BSA, was added to the tissue sections and incubated overnight at 4°C on an orbital shaker. The next day, tissue sections were allowed to equilibrate at room temperature for 1 hour on an orbital shaker. Afterward, tissue sections were washed three times with PBS containing 0.1% Triton X-100. Goat anti-Rabbit IgG Alexa Fluor Plus 488 highly absorbed (Thermo Scientific #A32731) or goat anti-mouse IgG Alexa Fluor Plus 555 highly absorbed (Thermo Scientific #A32727) secondary antibodies, diluted in PBS with 1% BSA, were added to the tissue sections and incubated for 1 hour at room temperature on an orbital shaker. Subsequently, tissue sections were washed twice with PBS containing 0.1% Triton X-100 for 5 minutes each, followed by two washes with PBS for 10 minutes each. The tissue sections were then mounted on glass slides, allowed to air-dry, and a hydrophobic barrier was drawn around the tissue. If dual staining was not performed, NeuroTrace Red (Thermo Scientific #N21482), diluted 1:100 in PBS, was added to the sections and incubated for 30 minutes on an orbital shaker. After incubation, tissue sections were washed three times with PBS for 10 minutes each. TrueBlack (Biotium #23007) was applied for 10 minutes and the tissue sections were coverslipped using ProLong Antifade mounting medium containing DAPI (Thermo Scientific #P36935) to counterstain nuclei. The stained sections were then allowed to dry completely before microscopic examination on BZ-X All-in-One Fluorescence microscope (Keyence, Itasca, IL).

### Regional immunoreactivity strength score for JNK3

Eight matching sections were selected from each animal and subjected to immunohistochemical staining for JNK3 (DAB) as described previously [50]. Utilizing a BZ-X All-in-One microscope (Keyence, Itasca, IL), a single observer scored the immunoreactivity strength to JNK3. The scoring rubric applied ranged from 0-3: 0 signified the absence of reactivity, 1 represented mild reactivity, 2 corresponded to moderate reactivity, and 3 indicated high reactivity. A visual representation of this scoring method can be viewed in Figure 3. An average of 3 animals for each sex was take for regions quantified. To provide comprehensive, visual representations, spatial mappings were created with Adobe Photoshop 2023. These mappings, presented as a six-color gradient scale, encompass eight brain atlas sections [50]. The sections were adapted from the Allen Mouse Brain Atlas (Allen Institute of Brain Science, 2011) and Paxinos and Franklin’s “The Mouse Brain in Stereotaxic Coordinates,” Fifth Edition [50].

### Western Blotting

Fluorescent semiquantitative western blotting was performed on brain cerebellum, brainstem, hippocampus, midbrain, striatum and cortex obtained from 12-month-old male and female C57BL/6J mice. The brain tissues were homogenized using N-PER lysis buffer (Thermo Scientific #87792) supplemented with protease and phosphatase inhibitors without EDTA (Thermo Scientific #1861281). Protein concentration was determined by performing a bicinchoninic acid (BCA) assay (Thermo Scientific #23225) following the manufacturer’s protocol. Equal amounts of protein (20μg) samples were then subjected to SDS-PAGE. Proteins were transferred onto nitrocellulose membranes using a BioRad TurboTransfer system. The membranes were blocked with 5% non-fat milk in Tris-buffered saline (TBS) for 1 hour at room temperature rocking. Primary antibodies (Table 1), diluted in TBS containing 0.1% Tween20 (TBST), were added to the membranes and incubated overnight at 4°C rocking. Following primary antibody incubation, membranes were washed three times for 7 minutes each with TBST. Anti-rabbit IgG (Cell Signaling #5151S) and anti-mouse IgG (Cell Signaling #5470S) fluorophore-conjugated secondary antibodies were then applied to the membranes and incubated for 1 hour at room temperature rocking. After secondary antibody incubation, membranes were washed three times for 7 minutes each with TBST. Fluorescent signals were detected and captured using Odyssey CLx Imaging System. Images of the western blots were imported directly into the Empiria Software (LI-COR Biosciences, Lincoln, NE). Subsequently, these images were quantified in relation to housekeeping gene expression or total protein stain.

### Phos-tag Western Blotting

For Phos-tag Western blotting, equal amounts (20μg) of previously prepared brain lysate samples were loaded into SuperSep Phos-tag pre-cast gels (FUJIFILM Wako Chemicals # 198-17981) and subjected to electrophoresis. Following electrophoresis, the gels were washed three times for 20 minutes each with transfer buffer containing 20% methanol, 10% stock transfer buffer (30g TrisBase and 144g Glycine in 1 littler deionized water), 70% deionized water, and 10 mM EDTA. The gels underwent a final wash with transfer buffer to eliminate any residual EDTA. Next, proteins were transferred from the gel to a nitrocellulose membrane using a conventional wet transfer method as described by SuperSep Phos-tag precasted gels. The membrane was then washed with TBS for 10 minutes. This was followed by drying the membrane in 37°C bench incubator for 10 minutes and subsequent rehydration using TBS. Total protein staining was then carried out on the membrane using the REVERT Total Protein Stain (LI-COR Biosciences) #926-11010 following the manufacturer’s instructions. The membrane was imaged using the Odyssey CLx Imaging System. The membrane was blocked using 5% non-fat milk in TBS for 1 hour rocking at room temperature. The membrane was then incubated with a JNK3 primary antibody, dissolved in TBST containing 5% BSA, and left overnight at 4°C rocking. The following day, the membrane was washed thrice with TBST for 7 minutes each. Secondary antibodies were added to the membrane and rocked for 1 hour at room temperature. Following the incubation, the membrane was washed again three times with 1× TBST for 7 minutes each. The membrane was then imaged again using the Odyssey CLx Imaging System. The images were quantified and normalized to the total protein stain using Empiria Studio Software to determine relative protein expression levels. Images of the Western blots were imported directly into the Empiria Software (LI-COR Biosciences, Lincoln, NE). Subsequently, these images were quantified in relation to total protein staining and fold-changes were obtained.

### Cell Culture

Cerebral tissues were collected from mixed-sex C57BL/6J pups on postnatal day (PND) 0–1 and dissected for primary cortical neuron culture preparation. Dissociation of cortical cells was achieved using 1mg/mL of papain and 0.5U/mL of dispase. Post-dissociation, cells were washed, triturated, pelleted, and subsequently resuspended in the culture medium. The resultant cell suspension was seeded onto poly-D-lysine-coated 6-well plates at an average density of 1×10^6^ cells per well. Cells were cultured in a neurobasal medium, enriched with B-27 plus, 50 IU of penicillin, 50mg/mL of streptomycin, and 2 mM of L-glutamine. Half of the medium was refreshed 24 hours post-isolation. After four days, half of the medium was substituted with a culture medium supplemented with 2% B-27 plus supplement and 5μM of cytosine β-d-arabinofuranoside (Ara-C; Sigma; #C1768). The culture medium was then changed every three days. For microglia and astrocytes culture, brains were also collected from mixed C57BL/6J pups on PND 0-3, and then homogenized in a solution of 0.25% trypsin with EDTA. The homogenized cell suspension was relocated to a T175 flask containing DMEM medium, supplemented with 10% FBS, 2mM l-glutamine, 1mM Na-Pyruvate, 100μM non-essential amino acids, and 1% penicillin and streptomycin. On day 7, the culture medium within the flask was replaced. By day 14, mature microglial cells were isolated from astrocytes using EasySep Mouse CD11b Positive Selection Kit II (Stemcell Technologies #18970A,) and subsequently seeded into p60 plates. The cells were harvested using a RIPA buffer containing protease and phosphate inhibitors. Lysates were left on ice for 30 minutes, homogenized, and centrifuged to gather the supernatant. All samples were stored at -80°C until further use.

### Statistical Analyses

For western blots, fold values which were obtained from Empiria Software were then incorporated into the GraphPad Prism 9 Software, where a 2-way ANOVA was executed to analyze differences related to age, brain region, and sex.

## Results

### JNK3 localization in adult male and female mouse brain

Immunohistochemical (IHC) staining was employed to visualize the relative abundance of JNK3 within different brain regions of 12-month-old male and female C57BL6/J mice using a knockout-validated isoform-selective antibody (Supplemental Figure 1). JNK3 was consistently identifiable across all brain regions in sagittal brain sections, using DAB IHC-staining, with notably enriched concentrations in the cortex and hippocampus (Figure 1). JNK3 abundance was clear in the primary somatosensory cortex (Figure 1A) and the primary motor cortex (Figure 1B) particularly. In Figure 1C, robust JNK3 staining was noticed in the hippocampal formation. Additional areas with enriched JNK3 localization included the piriform cortex (Figure 1D), the olfactory tubercle (Figure 1E), and the areas of the thalamus (Figure 1F). Within these images, higher magnification (20× and 60×) images illustrate that the JNK3 distribution appears to be predominantly in neurons in these regions (Figure 1).

**Figure 1.**
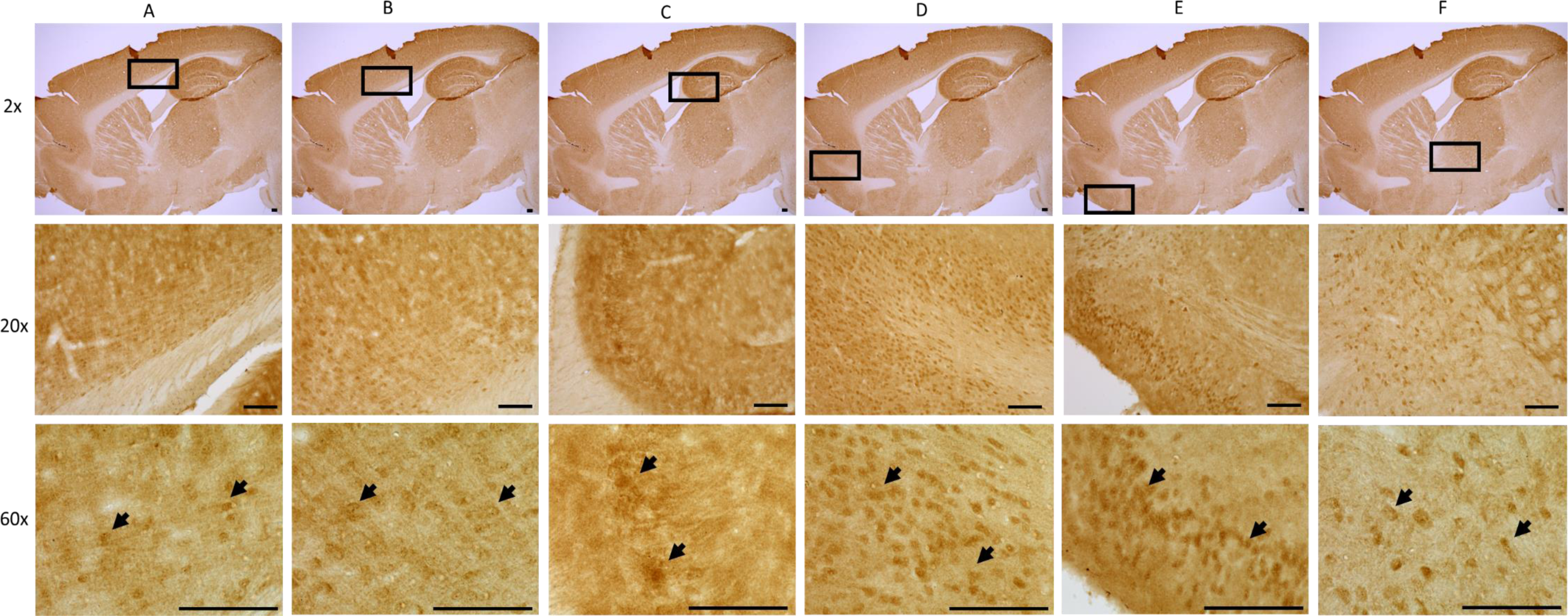
Localization of JNK3 in 12-month-old C57BL6/J male mice with immunohistochemical (DAB) staining. **A.** Primary somatosensory cortex. **B.** Primary motor cortex. **C.** CA3 of the hippocampus. **D.** Piriform cortex. **E.** Olfactory tubercle. **F.** Thalamic Region (Lateral 1.68mm). Increasing levels of magnification are indicated along the left side of the rows as 2×, 20×, and 60×. Scale bar = 100μm. The black boxes at 2× magnification highlight the areas of magnification for the 20× and 60× images for each column. Black arrows in the 60× panel identify neurons with JNK3 positive signal in the above indicated regions.

### JNK3 localization in the brainstem of adult mouse brain

IHC staining was employed on sagittal slices of C57BL6/J mouse brains to analyze JNK3 distribution in specific regions of the brainstem (Figure 2). JNK3 staining was observed throughout the brainstem, albeit more diffuse than in the cortical and hippocampal regions (Figure 1). The JNK3 distribution in the brainstem was consistent more with specific nuclei. Specifically, JNK3 prominently appears in the spinal trigeminal nucleus (Figure 2A), the intermediate reticular nucleus (Figure 2B), the facial nucleus (Figure 2C), the motor trigeminal nucleus (Figure 2D), the vestibular nucleus (Figure 2E), and the interposed cerebellar nucleus (Figure 2F). Significant staining was also observed in the cerebellar cortex of the anterior and posterior lobes (Figure 2F). As in other brain regions, JNK3 localization at higher magnification appears to be almost exclusively neuronal consistent with previous histological reports [7, 8].

**Figure 2.**
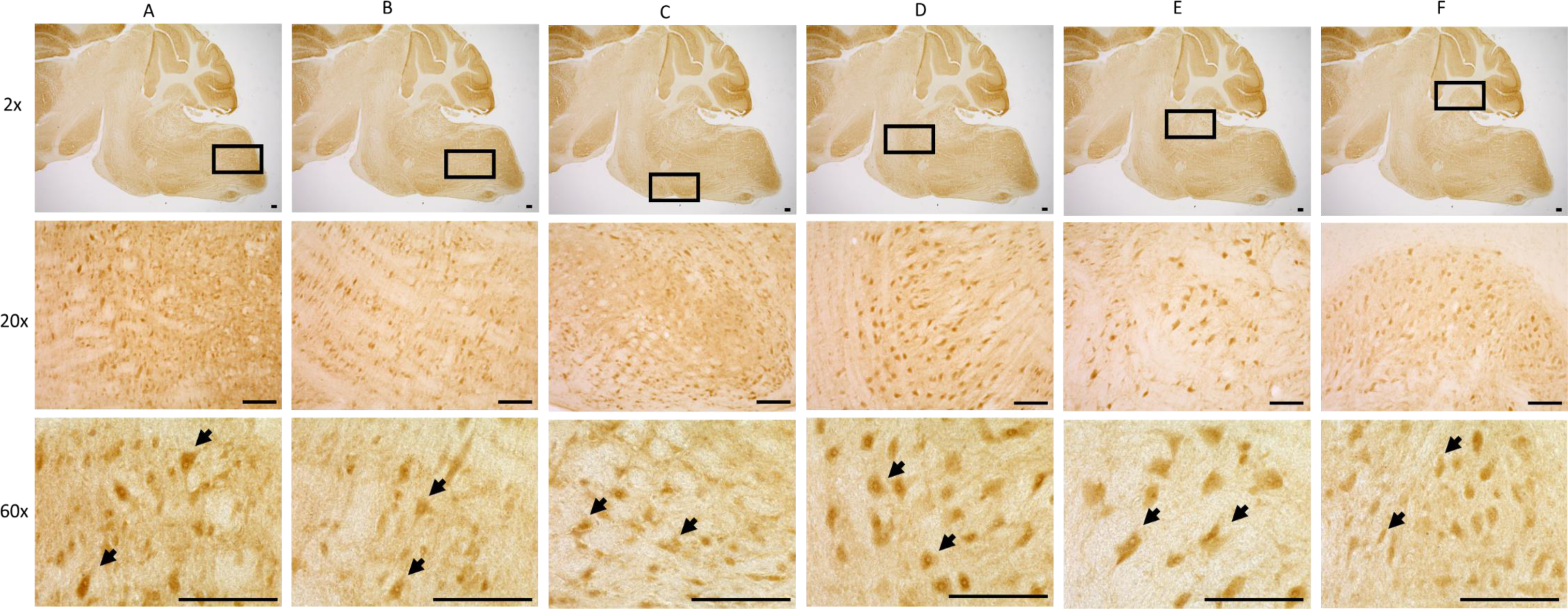
Localization of JNK3 in 12-month-old C57BL6/J male mice with immunohistochemical (DAB) staining. **A.** Spinal trigeminal nucleus. **B.** Intermediate reticular nucleus. **C.** Facial nucleus. **D.** Motor trigeminal nucleus. **E.** Vestibular nucleus. **F.** Interposed cerebellar nucleus. (Lateral 1.68mm) Scale bar = 100μm. The black boxes at 2× magnification highlight the areas of magnification for the 20× and 60× images for each column. Black arrows in the 60× panel identify neurons with JNK3 positive signal in the above indicated regions.

### Qualitative anaysis of JNK3 levels in the adult mouse brain and cervical spinal cord

To semi-quantify JNK3 abundance in each region, mouse brains were coronally sectioned and IHC-stained with DAB. Based on the relative intensities of JNK3 signal in these sections, JNK3 immunoreactivity was characterized as low (Figure 3A), moderate (Figure 3B), or high (Figure 3C). These benchmarks were then used to form a scored rubric of JNK3 abundance across the brain. From the anterior to the posterior, high JNK3 immunoreactivity was noted in the anterior cortical regions, including the agranular insular cortex and dorsal peduncular cortex (Table 2). Higher immunoreactivity was observed in the cingulate cortex, primary and secondary motor cortices, primary and secondary somatosensory cortices, and the piriform cortex (Table 2, Figure 4A-C). High JNK3 immunoreactivity was also evident in the olfactory tubercle (Table 2, Figure 4A-B). Contrariwise, the striatum and the globus pallidus in the same region exhibited low JNK3 immunoreactivity (Table 2, Figure 4A-C). The immunoreactivity for JNK3 was consistent throughout the striatum, with the internal globus pallidus demonstrating slightly higher immunoreactivity for JNK3 than the external globus pallidus (Figure 4C). Transitioning into the temporal lobe, the auditory cortex and other cortical regions showed high immunoreactivity for JNK3 (Table 2, Figure 4C-D). The hippocampus demonstrated significantly high JNK3 immunoreactivity, particularly in the CA1-CA3 regions and the dentate gyrus; whereas, the subiculum had very low JNK3 immunoreactivity (Table 2, Figure 4C-D). Near the hippocampus, the thalamus and hypothalamus were observable (Table 2, Figure 4C), both displaying moderate to high immunoreactivity for JNK3. Consistently, most staining in the thalamus was observed in the mediodorsal region, with slightly less in the ventral lateromedial region (Figure 4C). However, the amygdala, found within this region, had notably low immunoreactivity for JNK3 (Table 2, Figure 4D). The nearby habenular nucleus also showed considerable JNK3 immunoreactivity (Table 1, Figure 4C). As we moved closer to the brainstem, the cerebellar lobes and fissures were visible, although they displayed very low reactivity for JNK3. Within the anterior region of the brainstem, the midbrain, JNK3 exhibited high immunoreactivity in the cerebellar peduncle and motor trigeminal nucleus (Table 2; Figure 4E-F). JNK3 also showed high immunoreactivity in the olivary nucleus, pontine reticular nucleus, and raphe nucleus (Table 2; Figure 4E-F). Furthermore, JNK3 demonstrated moderate immunoreactivity in the cuneiform nucleus and low reactivity in the reticulotegmental nucleus (Table 2, Figure 4E-F). Transitioning into the posterior part of the brainstem, the medulla oblongata, JNK3 also demonstrated significantly high immunoreactivity in motor-associated areas, including the vagus nerve nucleus, hypoglossal nucleus, lateral reticular nucleus, and inferior olive nucleus (Table 2; Figure 4G). The cuneate nucleus also displayed high JNK3 immunoreactivity, albeit slightly less than the other regions (Table 2, Figure 4G). In the pons region of the brainstem, JNK3 displayed high immunoreactivity overall. This was particularly evident in the facial nucleus, cerebellar nucleus, vestibular nucleus, granular insular cortex, gigantocellular reticular nucleus, spinal trigeminal nucleus, and cochlear nucleus (Table 2, Figure 4F). The immunoreactivity observed in the cervical spine displayed marked similarity to that of the brainstem; wherein, JNK3 localization in the cervical spinal cord (Figure 4H) was also consistently high across nuclei in the grey matter but was absent in the lateral regions (anterior and posterior horns) and the funiculus (Figure 4H). Lastly, no significant sex differences were observed between male and female brains in terms of JNK3 distribution.

**Figure 3.**
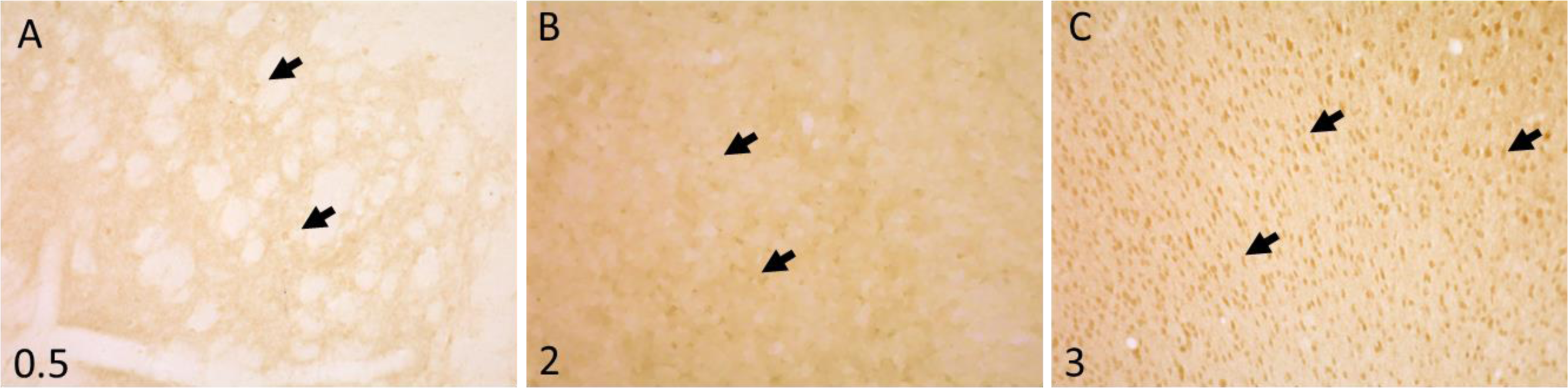
Photomicrographs of JNK3 immunoreactivity (DAB) in the brain of a C57BL/6J mouse categorizing the strength of reactivity: **A.** 0.5 indicates low reactivity **B.** 2 indicates moderate reactivity **C.** 3 indicates high reactivity. The black arrows indicate cells possessing the reactivity described in the qualitative rubric. Scale bar = 100μm.

**Table 2.**
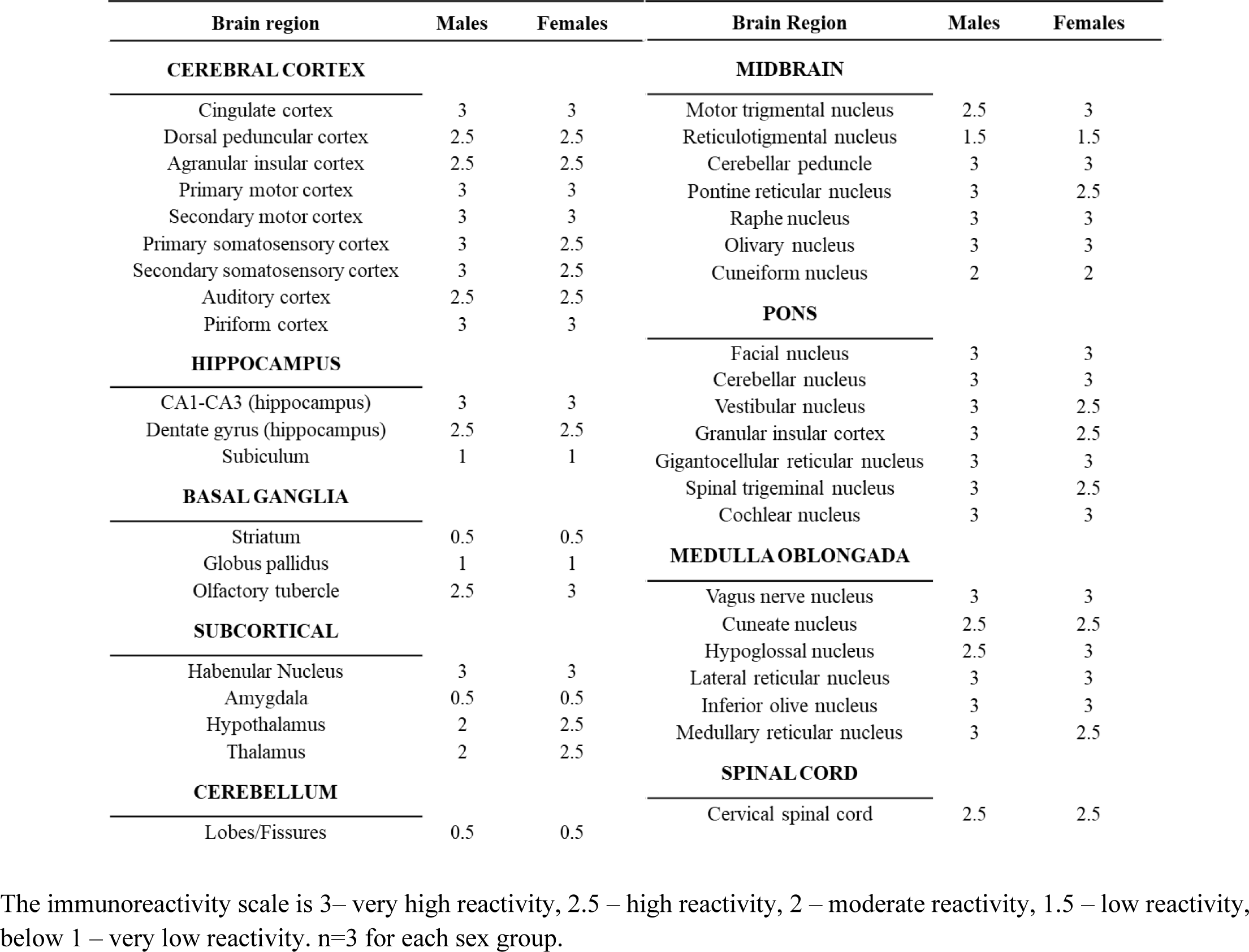
JNK3 immunoreactivity in the brains of C57BL/6J 12 months males and females.

**Figure 4.**
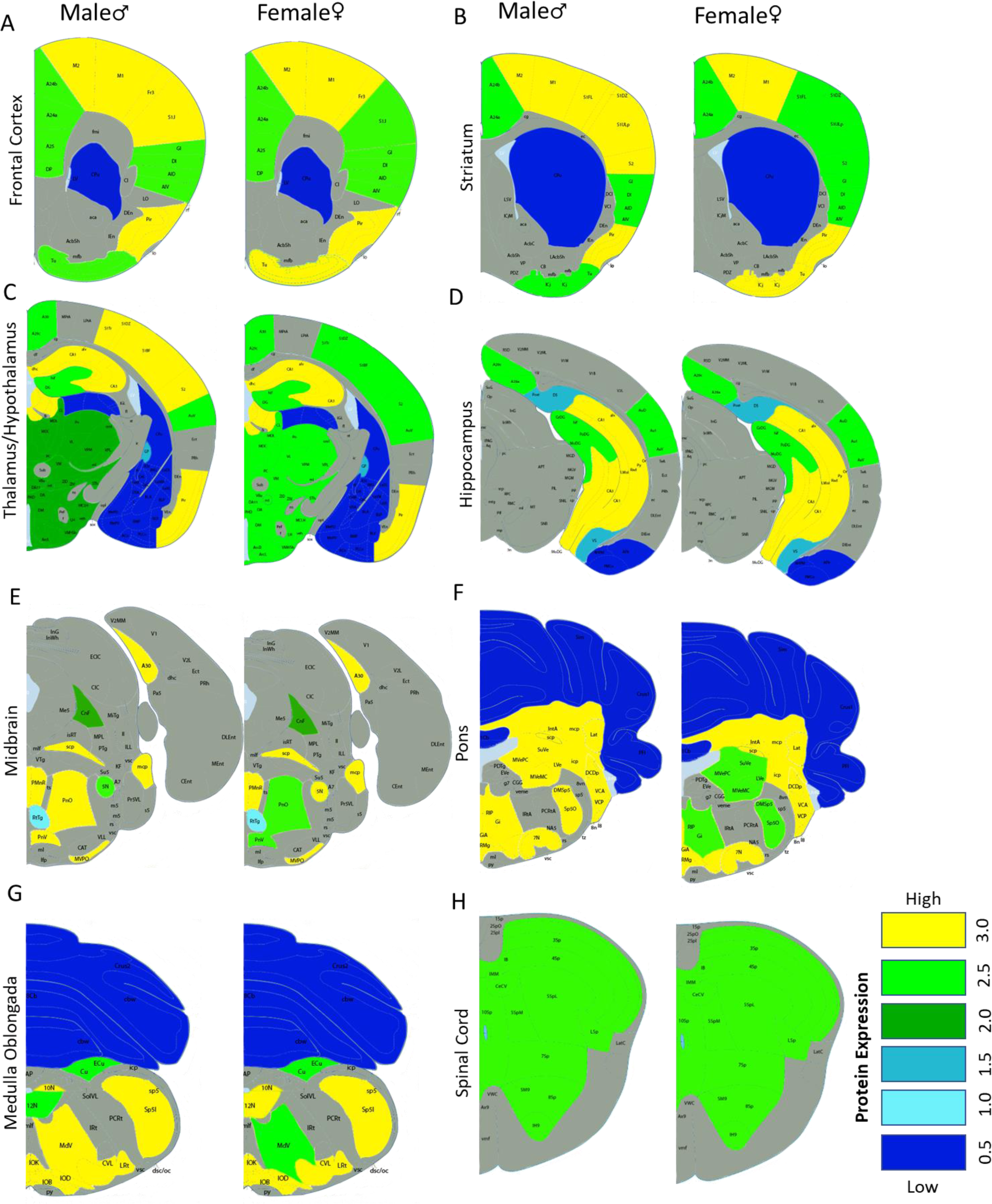
Spatial mapping of JNK3 immunoreactivity in the **A.** frontal cortex (bregma 1.33mm), **B.** striatal (bregma 0.73mm), **C.** thalamic (bregma -1.67mm), **D.** hippocampal (bregma -3.27mm), **E.** midbrain (bregma -4.83mm), **F.** pons (bregma -5.79mm), **G.** medulla oblongata (bregma - 7.19mm), and **H.** spinal cord (330 microns) regions. The expression of JNK3 is similar in males and females throughout the brain. The map was adapted from ***Table 1***. Gray areas were not quantified. The images and labeling were adapted from Alan Brain Atlas and Paxinos and Franklin’s The Mouse Brain Atlas.

### JNK3 co-localization with various neuron types

JNK3 distribution in our current results and in other histological studies has been described as primarily neuronal [7, 8]. To dissect the types of neurons that may express JNK3, co-immunofluorescent (co-IF) staining was performed with neuron-selective markers/antibodies and the JNK3 antibody. Across all regions of the C57BL6/J mouse brain, JNK3 exhibited extensive co-localization with NeuroTrace Red staining (Figure 5A). Antibodies for selective neuronal markers were used to discern the JNK3-positive neuron species. In the brainstem, an overlay of JNK3 and the motor neuronal marker, transcription factor islet 1 (Isl1), emerged throughout the region (Figure 5B). This co-localization was observed in areas critical for motor function, including the vestibular nuclei, pontine reticular nucleus, and all cranial nerve nuclei. Similar Isl1 localization also became apparent in the spinal cord, particularly in the anterior sections containing motor neurons. JNK3 also displayed co-localization with the cholinergic neuronal marker, choline acetyltransferase (ChAT), especially in regions densely populated with cholinergic neurons, such as the habenular nucleus (Figure 5C) and the cuneate nucleus in the brainstem. In dopaminergic neuron-rich regions, including the substantia nigra, co-localization of JNK3 with the dopaminergic neuronal marker, tyrosine hydroxylase (TH), also became evident (Figure 5D). The examination of JNK3 and glutamate decarboxylase (GAD67), a GABAergic marker, possessed overlapping signals that appeared across all brain regions, with higher intensity areas of the hippocampus and cortex shown in Figure 5E. In contrast, a lack of co-localization between JNK3 and the glutamatergic neuronal marker, vesicular glutamate transporter 1 (VGLUT1), was noted, even in areas known for a high concentration of glutamatergic neurons, such as the hippocampus (Figure 5E). These findings collectively suggest that JNK3 is localized in most types of neurons, but not all. The subcellular localization of JNK3 varied across different regions of the brain. Primarily, JNK3 appeared to be distributed throughout the entire cell body, whereas in the brainstem and spinal cord, JNK3 localization was more nuclear (Figure 5A).

**Figure 5.**
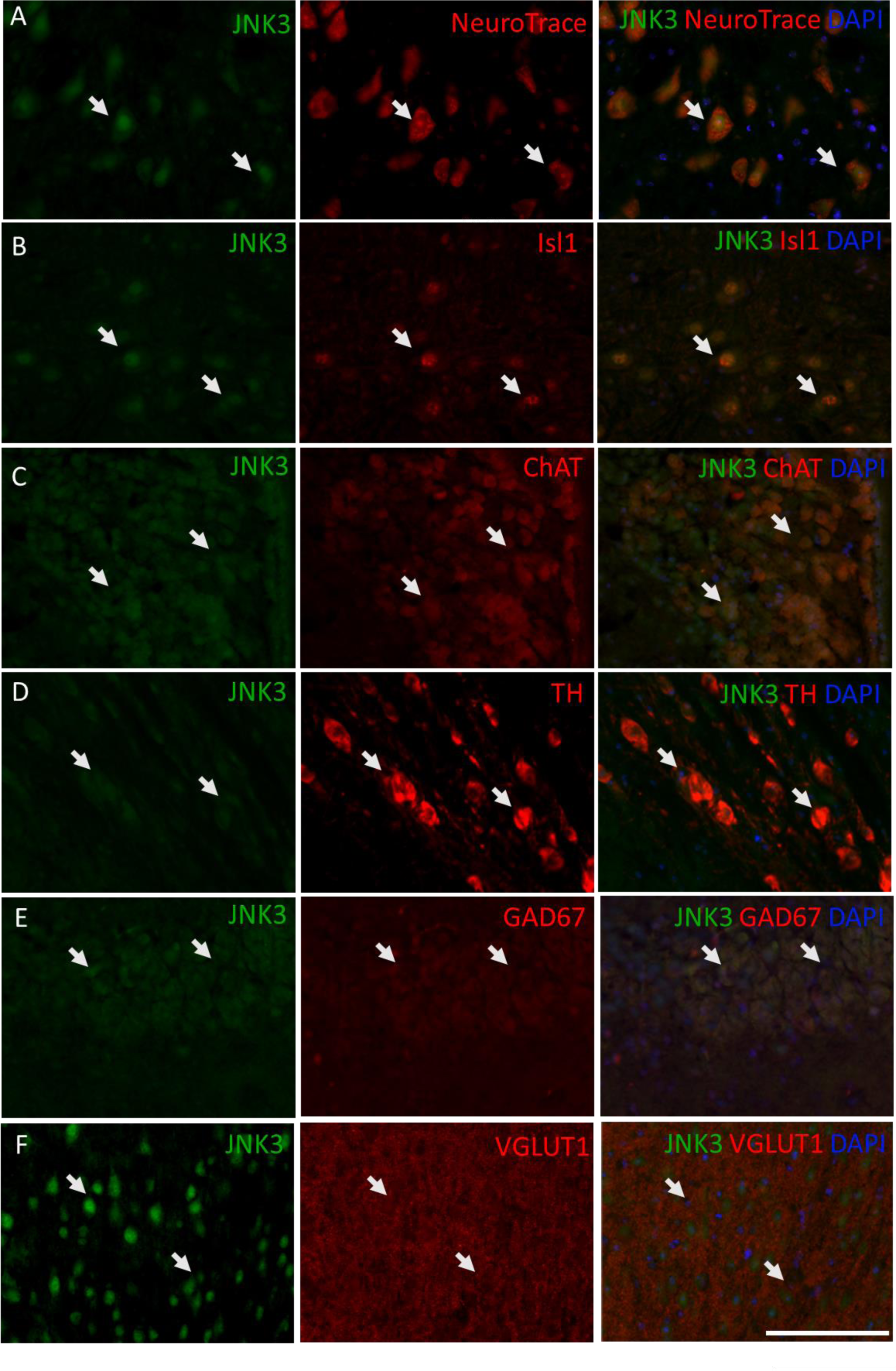
Co-localization of JNK3 with neurons in 12-month-old C57BL6/J male mice with immunofluorescent staining. **A.** JNK3 is present in neurons (NeuroTrace straining) in the facial nucleus. **B.** JNK3 co-localizes with a motor neuron marker (Isl1) in the hypoglossal nucleus. **C.** JNK3 co-localizes with the cholinergic neuron marker (ChAT) in primary cortex. **D.** JNK3 co-localizes with dopaminergic neuron marker (TH) in the substantia nigra. **E.** JNK3 co-localizes with GABAergic neuron marker (GAD67) in the CA3 of the hippocampus. **F.** JNK3 JNK3 does not co-localize with glutaminergic neuron marker (VGLUT1) in the primary cortex. White arrows indicate cells exhibiting colocalization of JNK3 and neuron-specific signals. Scale bar = 100μm.

### JNK3 protein levels in adult male and female brain

To corroborate the immunohistochemical and immunofluorescent findings, the protein levels across multiple regions of the C57BL6/J mouse brains were analyzed. Initially, a comparison of 3-month-old (the age of previous histological studies [7, 8]) and 12-month-old male mice revealed no significant differences in JNK3 levels were detected between the two age groups (Supplemental Figure 2). In 12-month-old mice, significantly higher JNK3 protein levels in the hippocampus and cortex compared to the cerebellum (Figure 6A). On the other hand, regions such as the brainstem, midbrain, and striatum exhibited JNK3 protein levels comparable to those in the cerebellum (Figure 6A). Semi-quantification of the protein levels was performed to illustrate any differences among the brain regions (Figure 6B). In addition, a sex-based comparison between male and female mice revealed no significant differences in JNK3 protein levels.

**Figure 6.**
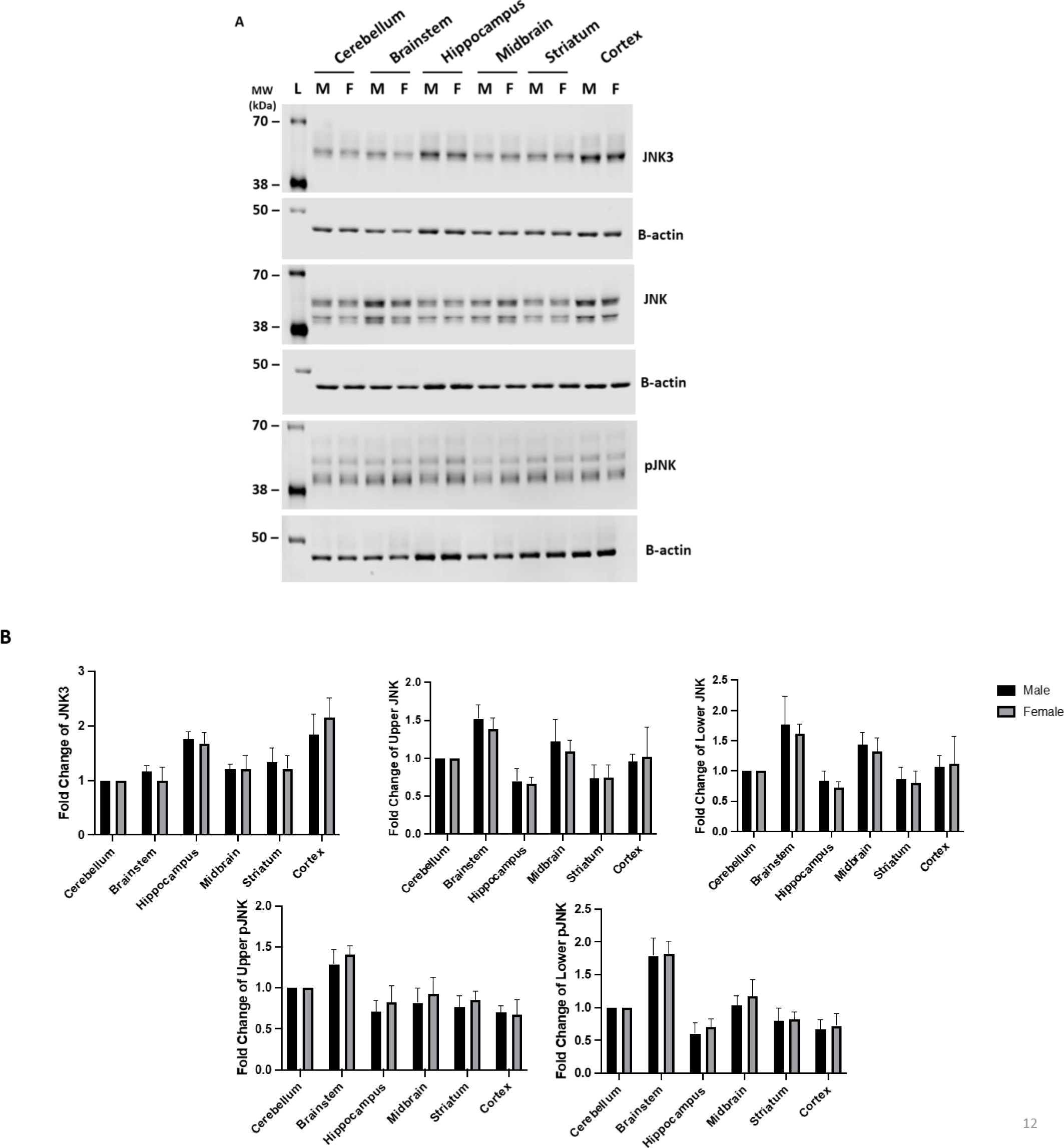
**A.** Sex comparison of JNK3, JNK, and pJNK protein levels in different brain regions of 12-month-old C57BL/6J mice. **B.** Analysis of JNK3, JNK, and pJNK protein levels comparing different brain regions of 12-month-old male and female C57BL/6J mice, with hippocampus and cortex being significantly higher in males and females. Brain regions were normalized to the cerebellum and quantified using Licor Empiria Software. Statistical analysis performed on Prism 9. Male n=5 and female n=6.

### Phosphorylated JNK and JNK3 abundance in adult male and female brain

Prevailing research suggests that elevated JNK3 activity may contribute to the induction of apoptosis; meanwhile, studies implicating of JNK3 in neuron morphology, neurotransmission, and behavior would indicate that elevated JNK3 activity in the healthy brain could illustrate areas that require JNK3 for normal neurophysiology. Thus, we used traditional and Phos-Tag© western blot analyses with total JNK antibodies and the JNK3-selective antibody to measure the abundance of JNK3 in dissected brain regions. A slight elevation of total JNK protein levels was observed in the brainstem and midbrain, while a decrease was observed in the hippocampus and striatum (Figure 7A). In contrast, the levels of phosphorylated JNK were mildly higher in the brainstem, while remaining consistent across all other regions (Figure 7A). Quantification of phosphorylated JNK3 in the brains of mice by Phos-Tag© western blotting revealed a higher proportion of phosphorylated JNK3 species in the hippocampus and cortex (Figure 7A & B). However, the brainstem presented fewer phosphorylated JNK3 species (Figure 7A & B). Upon further analysis of total JNK and phosphorylated JNK proteins, no statistically significant sex-based differences emerged between male and female mice regarding phosphorylated JNK, pJNK, and pJNK3 in the brain (Figure 7B). Thus, the presence of active JNK3 can be found at distinct concentrations within the brain.

**Figure 7.**
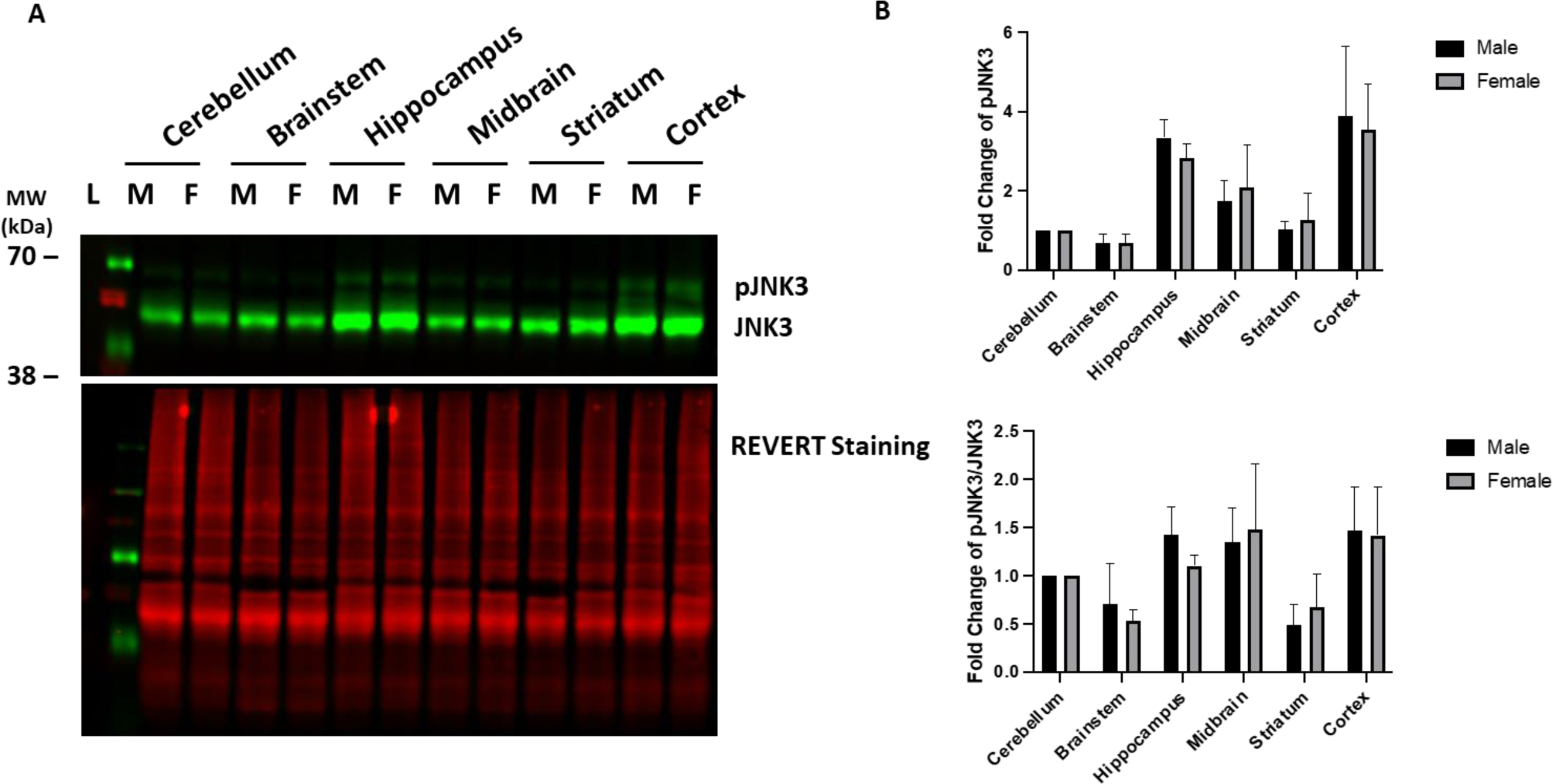
**A.** Sex comparison of pJNK3 protein levels in different brain regions of 12-month-old C57BL/6J mice. **B.** Analysis of pJNK3 protein levels comparing different brain regions of 12-month-old male and female C57BL/6J mice. Brain regions were normalized to the cerebellum and quantified using Empiria Software. Statistical analysis performed on Prism 9. Male n=4 and female n=4.

### JNK3 concentrations within astrocytes and in primary CNS cells

Recent studies have suggested that the expression of JNK3 may extend beyond neurons to glial cells, such as astrocytes and oligodendrocytes [33, 51–53]. To compare JNK3 abundance in other prominent cell types in the brain, primary neurons, astrocytes, and microglia were cultured from C57BL/6J mice. JNK3 was found in the cultured neurons and astrocytes (Figure 8A); meanwhile, JNK3 was not detectable in primary microglia (Figure 8A).Further examination of potential co-localization of JNK3 with astrocytes employed glial fibrillary acidic protein (GFAP) in IHC staining (Figure 8B). However, alignment of JNK3 staining with astrocyte localization did not occur in adjacent brain slices, indicating the absence of JNK3 in regions rich with astrocytes (Figure 8B). Cell morphologies in these regions are noted in Supplemental Figure 3 for GFAP-positive and JNK3-positive cells. This result may suggest that the prevailing JNK3 species in the healthy adult brain may be affiliated with neurons, as suggested by previous studies.

**Figure 8.**
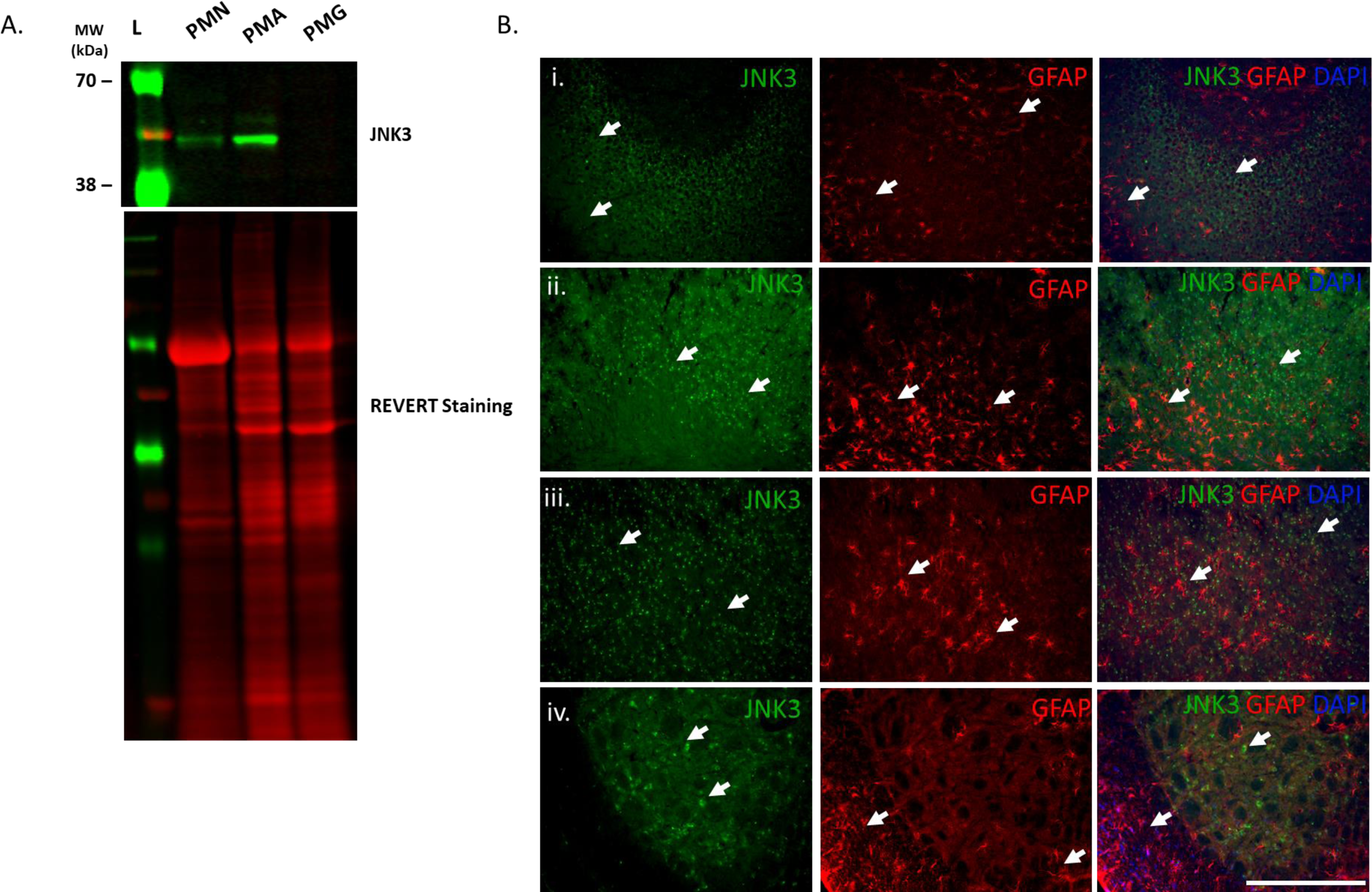
**A.** JNK3 abundance in primary cells from C57BL/6J mice. Cell lysates were normalized to REVERT staining and quantified using Licor Empiria Software. The molecular weight standards are indicated by L, primary mouse neurons are marked as PMN, Primary mouse astrocytes are indicated as PMA, and Primary mouse microglia are labeled as PMG. Statistical analysis performed on Prism 9. n=3. **B.** Localization of GFAP and JNK3 in adjacent coronal slices of C57BL6/J mice brain: **i.** CA3 of the hippocampus, **ii.** primary motor cortex, **iii.** forceps minor of the corpus callosum, and **iv.** spinal trigeminal region. The white arrows indicate the cells possessing either a JNK3 positive or GFAP positive signal. Scale bar = 100μm.

## Discussion

JNK3 is emerging as not only a central element of neurodegenerative disease pathophysiology, but also for relevance to normal neurological function. Our study builds upon previous JNK3 histological and neurological studies and provides a greater resolution of JNK3 distribution in the adult brain to help understand the kinase’s roles within the nervous system. The findings of our current study demonstrate that JNK3 is distributed throughout the brain in healthy adult C57BL6/J mice with protein (Figure 6) and phosphoprotein (Figure 7) levels elevated in the cortex and the hippocampus as observed in previous studies [7, 8, 48]. However, our study uncovers the enrichment of JNK3 and pJNK3 in areas of the brainstem, cerebellum, and cervical spinal cord not mentioned in previous histological investigations. Furthermore, the study reaffirms the predominance of JNK3 in neurons and identifies the specific types and locations of JNK3-positive neurons. Collectively, these data set the stage to examine the nuanced functions of JNK3 in these areas of the brain concerning motor function, cognitive abilities, disease progression, and other neurological processes, but the distribution of JNK3 from our studies may provide insights into the vulnerability of neurons in distinct brain regions to neurotoxic exposures and hallmark events in neurodegenerative diseases.

This study replicated earlier results with the p493F12 antibody, which indicated elevated JNK3 protein expression in the cortex and hippocampus, with diminished presence in the striatum [7, 8]. The cortical and hippocampal localization (Figure 1) corresponds with not only the proposed involvement of JNK3 in AD and related dementias (ADRD) [25, 26, 37, 38], but it also aligns with the putative involvement of JNK3 in neurobehavioral outcomes [46, 54]. As mentioned, earlier *Jnk3^-/-^* mice exhibit some behavioral deficits in the elevated plus maze and potentially in the Morris water maze compared to wild-type mice, and these tests require processes affiliated with the cortical and hippocampal regions suggesting that JNK3 activities may be crucial to basal neurological function [46]. Our current study finds that pJNK3 levels are elevated in the hippocampus and cortex of healthy male and female mice (Figure 7) indicating that there is basal JNK3 activities are contributing to neurophysiology in these regions, which is consistent with recent pJNK3 radioligand labeling studies [49]. The higher levels of JNK3 activity in the hippocampal region could also represent a “double-edged sword” to explain why these regions are vulnerable to neuron loss in AD and other neurodegenerative disorders, as elevated and sustained levels of pJNK3 have been implicated in these conditions. Perhaps, the physiological dependence on JNK3 in the cortex and hippocampus, as well as other areas with elevated pJNK3, may lower the threshold for triggering detrimental JNK3 signaling events. One possible mechanism for this might be the physiological phosphorylation of Bcl-2 and BH3-only protein members [55] that could lead to apoptotic priming and polarization of mitochondria and cellular pathways towards cell death [56]. Additionally, inhibition of mitochondrial JNK signaling in the brain is neuroprotective *in vitro* and *in vivo* in part by altering the phospho-status of Bcl-2 superfamily proteins [4, 57]. Indeed, the mitochondrial scaffold for JNK, SH3-binding protein 5 (SH3BP5 or Sab) is also highly expressed in the hippocampus and contributes to apoptosis and neurotransmission [58]. Thus, the tight coupling of JNK3 activity to basal neurophysiology in the hippocampus and other regions may be a vulnerability when neurotoxic insults arise.

Alternatively, the diminished presence of JNK3 in the striatum (Figure 1, Figure 4, and Table 2), a site of Parkinson’s disease pathology (the loss of dopaminergic terminals [59]), is interesting given not only the involvement of JNK3 in synapse dysfunction and dendritic pruning [15, 60], but also many data indicating that JNK3-selective inhibitors protect in part against the loss of striatal dopamine in preclinical models [11, 26, 49, 61]. The lower levels of JNK3 in the striatum may speak to a reduced capacity for neuritogenesis. The first description of JNK3’s role in neurite initiation was demonstrated when *Jnk3^-/-^*mice and JNK3-selective inhibitors prevented neuritogenesis of dorsal root ganglion neurons [62]; JNK3 was later implicated in neurite initiation in other neurons, including dopaminergic neurons [63, 64]. Our analyses of pJNK3 levels in striatum did not reveal robust levels of active JNK3, but if the pJNK3 species are restricted to synaptic terminals or neurites in this large area, the pJNK3 could be diluted from highly concentrated, but small regions of the area. Another possibility is that JNK3 could be signaling from dopaminergic soma in the substantia nigra, as midbrain levels of pJNK3 were found in male and female adult mice (Figure 7), and JNK3 protein levels were high in the substantia nigra (Figure 4) as well. Further, Bales, et al. found that in a preclinical model of PD, pJNK3 levels significantly increased in the midbrain and in the area of the substantia nigra [49]. These data indicate that the increase in JNK3 activity for extended duration may be neurodegenerative [26], and the ability to detect these increased may be a useful way to identify vulnerable brain regions or to monitor disease progression to better inform treatment of individual patients [49].

Intriguingly, our study indicates that there are moderate JNK3 protein levels in various subcortical areas, including the thalamus and hypothalamus (Figures 1 & 4 and Table 2). The recent implications of JNK3 in metabolic homeostasis and disease [47, 65, 66] illustrate its possible function in these regions. Specifically, smaller neurons of the supraoptic nucleus exhibited an modest increase in JNK3 following tetrodotoxin treatment *in vitro* [65]. Additionally, brain-penetrant JNK inhibitors with partial selectivity for JNK3, namely SR11935, reduced food intake and body mass in adult mice [67]. It appears that JNK3/JNK2 inhibition sensitized animals to leptin effects by increasing leptin-mediated activation of signal transducer and activator of transcription 3 (STAT3) through the downregulation of suppressor of cytokine signaling 3 (SOCS3) [67]. Thus, while there may be therapeutic advantages to targeting JNK3 in neurological disorders, the specificity (i.e., targeting mitochondrial JNK signaling) and dose of these prospective interventions will require careful consideration to prevent effects on basal neurological functions of JNK3.

The distribution of JNK3 was found to span the entire brainstem and extend into the cervical spinal cord (Figure 2 & 4 and Table 2). Parallel findings were made previously in AD patient brains; wherein, JNK3 localization was noted not just in the cortical and hippocampal regions, but also within the brainstem and spinal cord [8]. Intriguingly, JNK3 colocalized with the motor neuron marker Isl1 in the brainstem and spinal cord with a prevailing nuclear appearance in its subcellular localization (Figure 5). The presence of JNK3 in motor neurons fits with the identification of JNK3 in other brain regions implicated in motor functions, such as the motor cortices, basal ganglia, cerebellum, and brainstem nuclei (Figure 1, 2, and 4). Examining pJNK3 levels in these regions, especially the cortices, cerebellum, and brainstem indicate there is active JNK3 in these regions with higher levels of activity in the cortex and cerebellum when compared to the brainstem. Nonetheless, these observations may indicate that JNK3 may have a role in the regulation of motor functions. Past research has implicated JNK3 activity in neurodegenerative conditions associated with motor neurons, such as spinal cord injury, spinal muscular atrophy (SMA) and amyotrophic lateral sclerosis (ALS) [68–70]. These studies suggest that elevated pJNK3 initiates motor neuron loss and that inhibition of JNK3 by small molecules or gene ablation can protect motor neurons [68–71]. The presence of JNK3 and pJNK3 in motor regions is an noteworthy result given the absence of apparent motor defects in knockout animals or those treated with JNK3-selective inhibitors [5]; nonetheless, the magnitude of JNK3 levels may be indicative of an inherent vulnerability of motor neurons in these regions to stress.

Prior research has identified JNK3 as primarily neuronal, specifically apparent in pyramidal neurons located in the hippocampus [8, 72], yet many JNK3 CNS activities highlighted here span across distinct neuron types. Moreover, the co-localization of JNK3 with specific neuronal types has been scarcely documented. Our study reports the co-localization of JNK3 with a variety of neuronal types, including motor neurons and dopaminergic neurons mentioned earlier, and cholinergic and GABAergic neurons. In AD, the loss of cholinergic neurons precipitates disease progression and may coincide with increases in pJNK3 along the course of disease [73]. Similarly, the role for JNK3 in the maintenance and survival of GABAergic neurons may align with alterations in GABAergic signaling that might contribute to the cognitive impairments seen in AD [74]. A particularly interesting results was the absence of co-localization between JNK3 and VGLUT1 because of JNK3’s relationships with endosomes and vesicular trafficking [41–44]. Postulation on this result may explain why JNK3 could be a crucial inducer of glutamate excitotoxicity in non-glutamatergic neurons, and that its minimal expression in VGLUT1-positive neurons could provide a level of protection. Cell-type-specific studies into the roles of JNK3 in distinct neurons could provide useful insights into its physiological and pathological activities in the brain and spinal cord.

In addition to neurons, astrocytes and oligodendrocytes have been identified as cell types that may have JNK3 expression within the CNS [52, 53, 75–77]. Our analyses of GFAP in adult mouse brain section do not align with JNK3 signals. However, primary cultured astrocytes do have substantial JNK3 concentrations consistent with previous studies implicating JNK3 in cultured astrocytes [52]. It is important to consider that our studies have been conducted on healthy adult brains, yet neurodegenerative conditions widely exhibit neuroinflammation and reactive gliosis [78] that could change the context of astrocytes allowing for JNK3 expression. Perhaps, these stresses may be similar to some of those in cell culture settings that trigger JNK3 expression in primary astrocytes. Unfortunately, we were unable to examine whether oligodendrocytes in the brain tissue possess JNK3 and pJNK3, but the relationship between oligodendrocytes and JNK3-implicated conditions like multiple sclerosis, suggests a potential interplay may exist. It is interesting that two CNS-resident glial cells (astrocytes and oligodendrocytes) closely related to neuronal function and neurotransmission would express JNK3. Studies are ongoing to determine if JNK3 glial activities may contribute to neurophysiology and disease progression.

In summary, JNK3 is broadly localized throughout a healthy adult mouse brain, with certain regions, such as the hippocampus and the cortex, displaying higher levels of JNK3 compared to others like the striatum. JNK3 is found in various neuronal cell types, including motor neurons, cholinergic neurons, dopaminergic neurons, and GABAergic neurons, but it appears to be absent from VGLUT1-positivve glutamatergic neurons. The activity of JNK3 is heightened in regions that exhibit high immunoreactivity for JNK3, such as the cortex and the hippocampus. In cell culture, JNK3 is present in neurons and astrocytes, though interestingly, in the brain, JNK3 does not co-localize with astrocytes. Overall, the insights gleaned from this study offer a higher resolution of the distribution of JNK3 and pJNK3 activity in the adult mouse brain, connecting long-standing observation in the field of neuro-JNK research. Given the inherent differences between mice and human neuroanatomy and CNS physiology, future studies will focus on conducting similar studies in human brain sections to define JNK3 and pJNK3 localization and subcellular distributions. Consequently, once the locations of JNK3 and active JNK3 are elucidated, this information will deepen our understanding of the potential mechanisms and vulnerabilities in neurodegenerative conditions, potentially setting the stage for the development of more effective diagnostic, prognostic, and/or therapeutic approaches.

## Acknowledgements

The authors would like to thank the members of the Chambers and Richardson labs for their helpful comments during the experiments and preparation of this manuscript. We are especially grateful to Dr. Melissa Edler (Kent State University) for her advice and encouragement during the imaging quantification and analyses from our study. We would like to thank Dr. Roger J. Davis (University of Massachusetts Chan Medical School and Howard Hughes Medical Institute) for the kind gift of the JNK-null MEFs we used to validate the JNK3 antibody for this study. Also, I would like to thank our Drs. Kim Tieu, Fenfei Leng, Timothy Allen, Hongxia Zhou, and Qin Li Cao for their helpful comments in the editing of the manuscript. This research was made possible by start-up funds provided by FIU to J.W.C and a grant from the Michael J. Fox Foundation for Parkinson’s Disease Research (#1211), and V.G. was supported by a research assistantship from the Transdisciplinary Biomolecular and Biomedical Sciences Training Program (NIH T32 GM132054-01). JRR was supported, in part, by NIH R01ES033892.

## Supplemental Figures and Legends

**Supplementa1 Figure 1.**
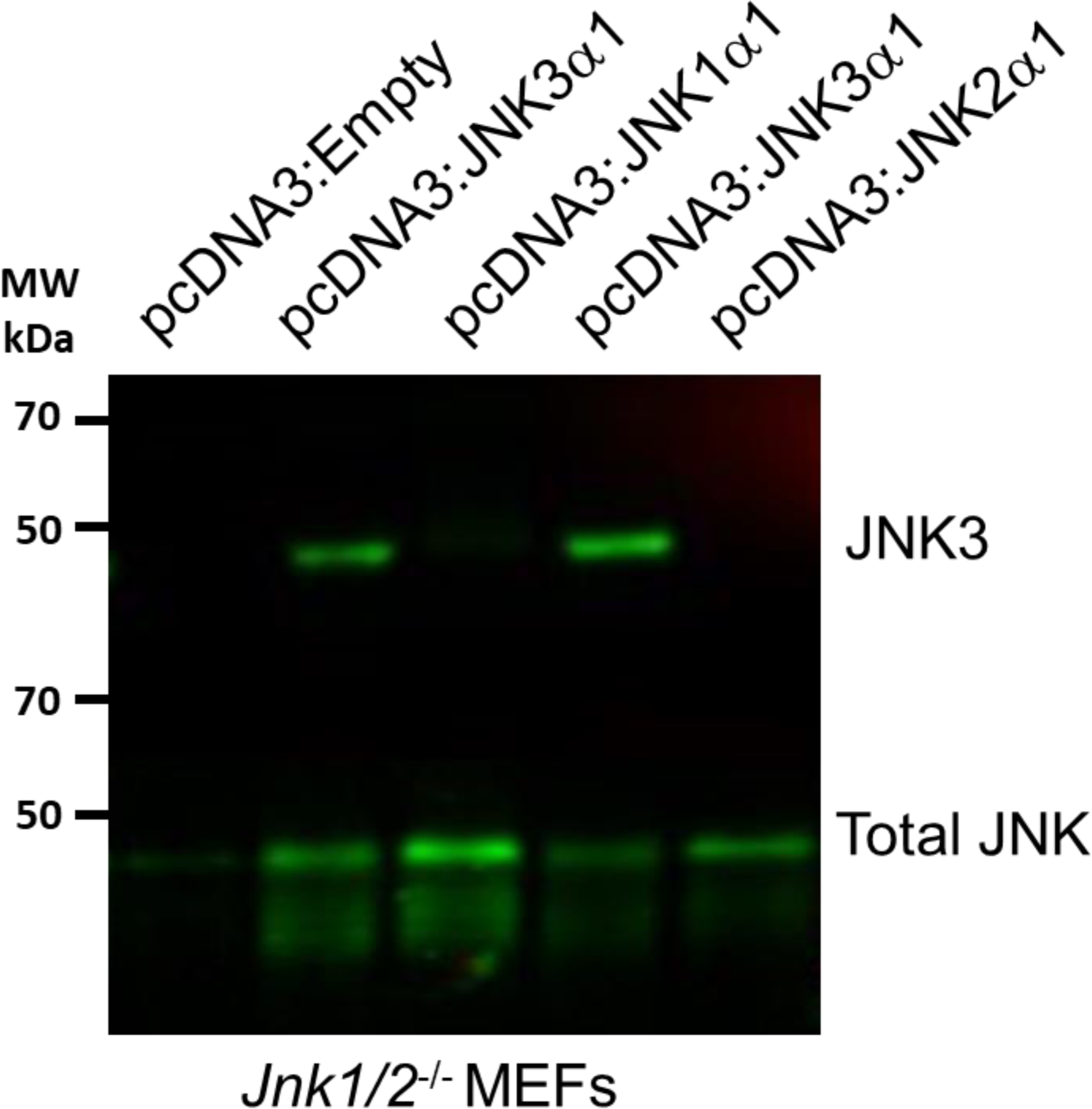
Validation of JNK3-selective antibody. JNK-null murine embryonic fibroblasts (a kind gift of Dr. Roger J. Davis) were transfected with empty vector or pCDNA encoding human JNK1α1, JNK2α1, and JNK3α1 variants. Cell lysates were then subjected to western blot analyses with JNK3-selective antibody (Cell Signaling) and Total JNK antibody (Cell Signaling).

**Supplementa1 Figure 2.**
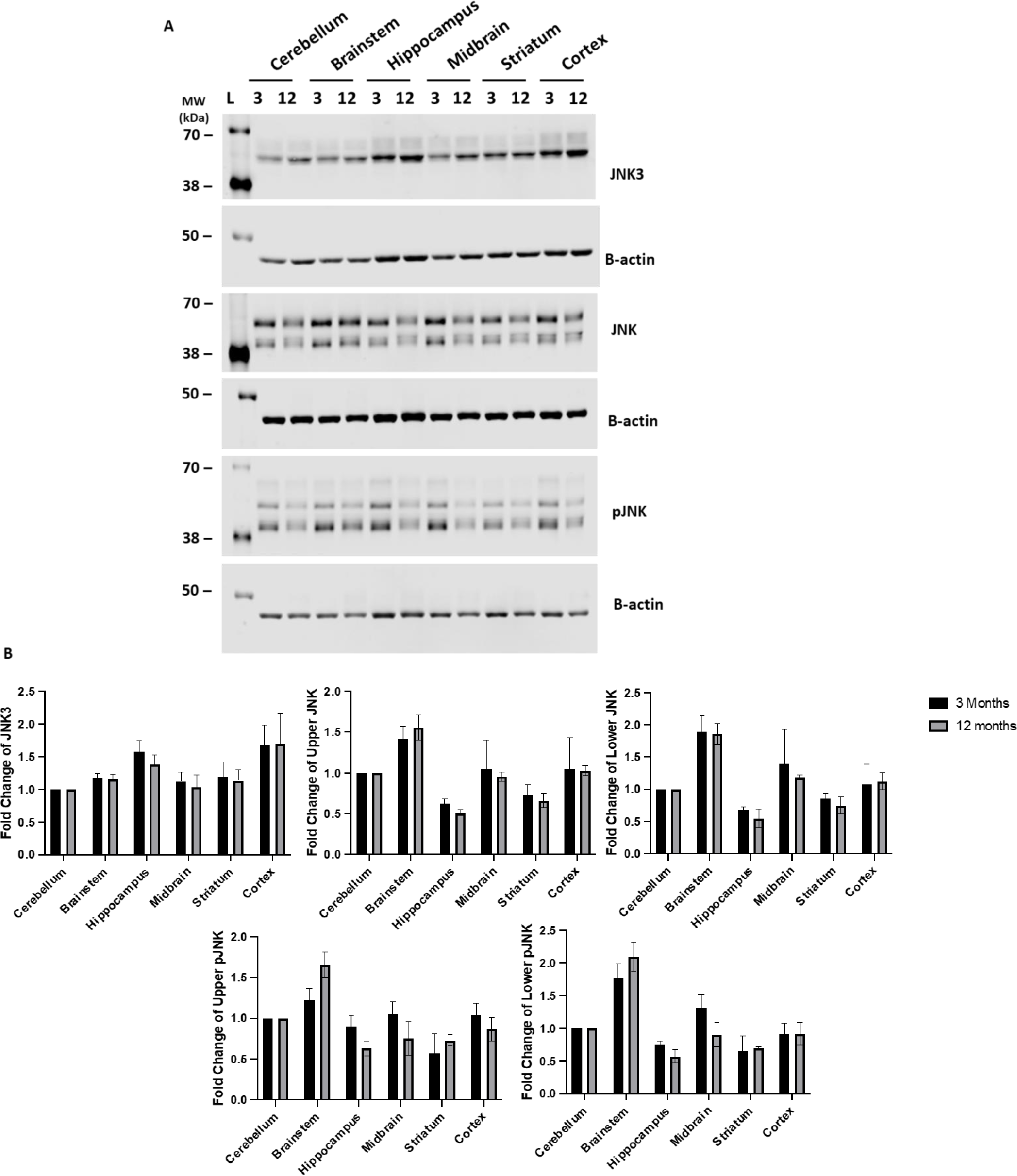
A. Age comparison of JNK3, JNK, and pJNK protein levels in different brain regions of 3 and 12 months old C57BL/6J mice. B. Analysis of JNK3, JNK, and pJNK protein levels comparing different brain regions of 3 and 12 months old male C57BL/6J mice. Brain regions were normalized to cerebellum and quantified using Empiria Software. n=3

**Supplementary Figure 3.**
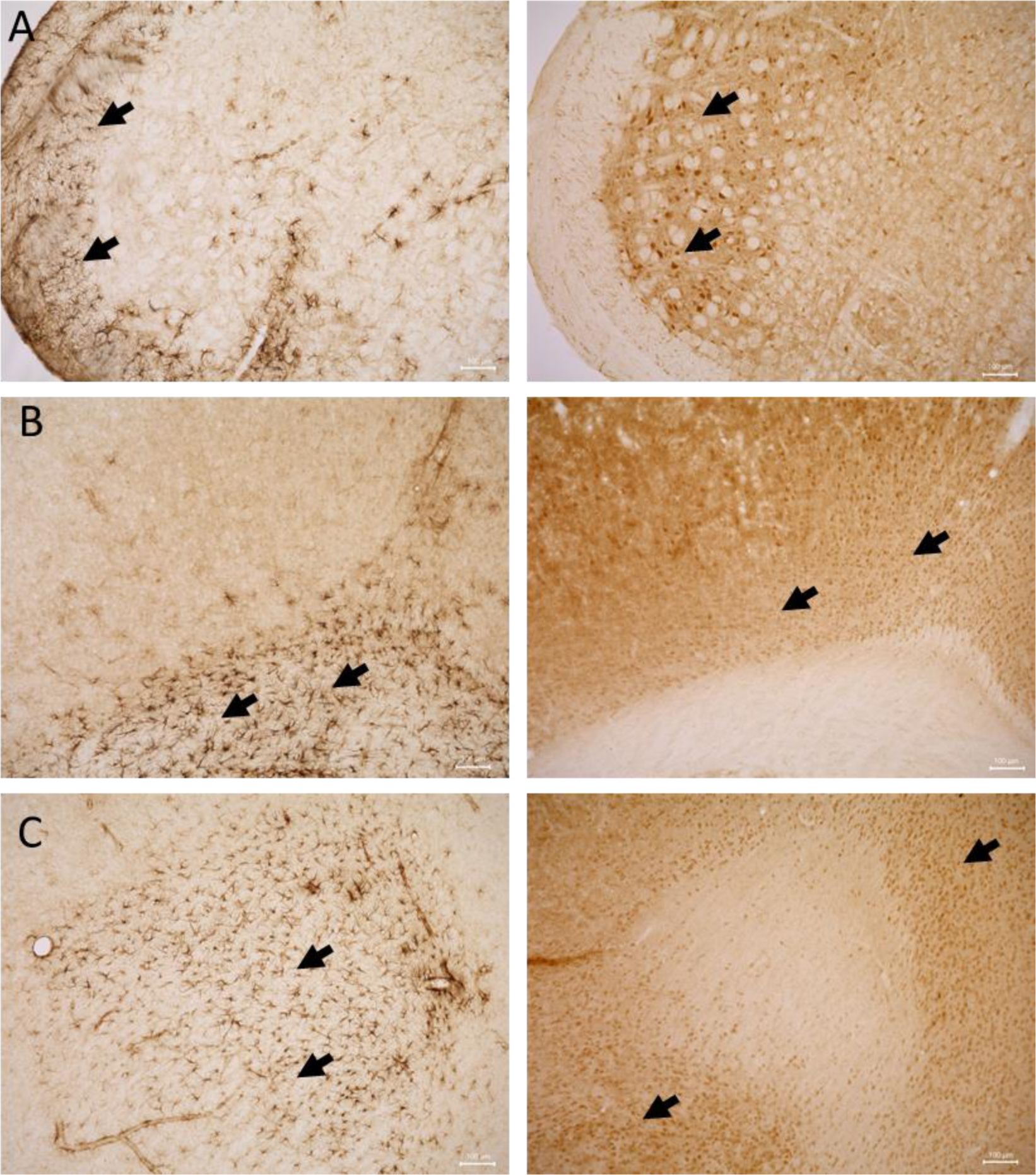
Localization of GFAP and JNK3 in adjacent coronal slices of C57BL6/J mice brain. (A) Localization of GFAP and JNK3 in the spinal trigeminal region (B) Localization of GFAP and JNK3 in the primary motor cortex. (C) Localization of GFAP and JNK3 in the forceps minor of the corpus callosum. Black arrows identify representative cells with the respective proteins present. Scale bar = 100 μm.

